# Loss of SPECC1L in cranial neural crest cells results in increased hedgehog signaling and frontonasal dysplasia

**DOI:** 10.1101/2025.11.21.689834

**Authors:** An J. Tran, Brittany M. Hufft-Martinez, Dana N. Thalman, Lorena Maili, Sean McKinney, Jeremy P. Goering, Paul A. Trainor, Irfan Saadi

## Abstract

SPECC1L encodes a cytoskeletal scaffolding protein that interacts with filamentous actin, microtubules, and cell junctional components. In humans, autosomal dominant mutations in *SPECC1L* cause a syndrome characterized by craniofrontonasal anomalies including broad nasal bridge, ocular hypertelorism, prominent forehead, and cleft lip/palate. Complete loss of *SPECC1L* in mice on a homogenous genetic background results in perinatal lethality, accompanied by subtle cranial differences and incompletely penetrant cleft palate. This lethality limits postnatal analysis of craniofacial development. Because cranial neural crest cells (CNCCs) contribute extensively to the formation of anterior craniofacial structures, we investigated whether disruption of SPECC1L in CNCCs contributes to the craniofrontonasal phenotypes observed in *SPECC1L*-related syndrome. We generated a *Specc1l*-floxed allele and crossed it with the *Wnt1-Cre2* deleter strain, which drives Cre recombinase expression in the dorsal neuroectoderm and NCCs. Most homozygous mutant *Specc1l*^*ΔCNCC*^ mutants survived postnatally and exhibited hallmark features of the human *SPECC1L*-related syndrome, including shortened skulls, reduced frontal bone area, nasal defects, and midface hypoplasia. The cranial mesenchyme of *Specc1l*^*ΔCNCC*^ mice displayed shortened primary cilia and increased Hedgehog (Hh) signaling activity at E13.5, as evidenced by enhanced GLI1 immunostaining. These defects were also observed early in E9.5 facial prominences, indicating that they are etiologic in nature. Collectively, *Specc1l*^*ΔCNCC*^ mice provide a novel model for investigating the roles of CNCCs, primary cilia, and Hh signaling in frontonasal prominence and midfacial development.

## Introduction

Craniofacial development is a highly coordinated process that depends on precise growth, migration, and differentiation of progenitor cell populations within embryonic facial processes.Central to this process are cranial neural crest cells (CNCCs), which populate the facial processes and give rise to most cranial and frontonasal structures (Jeong et al., 2004; Trainor, 2005; Tobin et al., 2008; Cordero et al., 2011; Sandell et al., 2011; Achilleos et al., 2015). Disruptions to this tightly regulated program often lead to craniofacial malformations, many of which stem from defects in key developmental signaling pathways. Among these, Hedgehog (Hh) signaling plays a central role in orchestrating facial morphogenesis. Excessive, insufficient, or otherwise dysregulated Hh activity can perturb the patterning and outgrowth of craniofacial structures, contributing to broad spectrum of congenital abnormalities ((Xu et al., 2023). Cytoskeletal scaffolding proteins play key roles in coordinating these events, integrating mechanical structure with intracellular signaling to guide tissue organization (Chiang et al., 1996; Mansouri et al., 1996; Bisgrove and Yost, 2006; Kuriyama and Mayor, 2008; Pollard and Cooper, 2009; Chang et al., 2016; Assis et al., 2017; Bleicher et al., 2020; Parker et al., 2020). Among them, SPECC1L (sperm antigen with calponin homology and coiled-coil domains 1-like) is essential for organizing the actin cytoskeleton, microtubules, and adherens junctions, thereby maintaining cellular integrity and enabling the coordinated growth required for craniofacial patterning (Saadi et al., 2011; Wilson et al., 2016; Hall et al., 2020; Goering et al., 2021a; Goering et al., 2021b; Mehta et al., 2023; Saadi et al., 2023).

In humans, autosomal dominant mutations in *SPECC1L* results in a spectrum of congenital craniofacial anomalies. Affected individuals commonly present with hypertelorism, broad nasal bridge, and cleft lip and/or palate, which are features consistent with frontonasal dysplasia group of disorders (Sedano and Gorlin, 1988; Farlie et al., 2016). Thus, *SPECC1L*-related hypertelorism syndrome is now considered as Teebi hypertelorism syndrome 1 (TBHS1) or brachycephalofrontonasal dysplasia (OMIM: 145420; ORPHA:1519), where the skull is short and wide. In addition to frontonasal dysplasia, patients can also manifest omphalocele, ear pits, uterine malformation, diaphragmatic hernia and congenital heart disease (Bhoj et al., 2015; Kruszka et al., 2015; Bhoj et al., 2019; Saadi et al., 2023).

The *SPECC1L* gene encodes a cytoskeletal scaffolding protein that has been shown to associate with microtubules, filamentous actin (F-actin), membrane-bound b-catenin, and non-muscle myosin II (Saadi et al., 2011; Wilson et al., 2016; Hall et al., 2020; Goering et al., 2021b). Loss of *Specc1l* resulted in perinatal lethality on both C57BL/6J and FVB/NJ backgrounds (Goering et al., 2021b). The homozygous null mutant embryos were frequently smaller overall and presented with subtle craniofacial anomalies. On the FVB/NJ background, the null mutants exhibited shortened primary cilia in the palate and ∼20% occurrence of cleft palate (Hufft-Martinez et al., 2025). Other *Specc1l* truncation and genetrap allele mutants displayed abnormally stabilized cell-cell adhesion between migratory (SOX10+) CNCCs (Wilson et al., 2016). Together, these findings suggest a role for SPECC1L in CNCC development and function.

To explore this role, we generated a *Specc1l* floxed allele and knocked out *Specc1l* in NCCs using the Wnt1-Cre2 driver line. Most of these conditional mutant mice survived postnatally and exhibited features consistent with the frontonasal dysplasia observed in patients with *SPECC1L-*related hypertelorism syndrome (Saadi et al., 2023), including altered skull length and width as well as frontal and parietal bone size. Mechanistically, cranial mesenchymal tissues from *Specc1l*^*ΔCNCC*^ embryos exhibited shortened primary cilia and elevated hedgehog (Hh) signaling activity, which is a critical regulator of midfacial growth (Chiang et al., 1996; Ahlgren and Bronner-Fraser, 1999; Bisgrove and Yost, 2006; Han et al., 2009; Goetz and Anderson, 2010; Briscoe and Thérond, 2013). These findings reveal a previously unrecognized role for SPECC1L in the cilia-mediated developmental signaling in CNCCs.

## Materials and Methods

### Mouse lines

The *Specc1l^fl^* allele was generated by CRISPR/Cas9-mediated recombination of loxP sites flanking exon 4 – the largest exon in *Specc1l.* We inserted the 5’ and 3’ loxP sites, sequentially, in mouse E14 embryonic stem cells (CVCL_C320), using the same CRISPR guide RNAs (gRNAs) that we used previously to generate the *Specc1l*^*ΔEx4*^ *null* allele (Goering et al., 2021b). The approximate genomic positions of the two gRNAs, 5′ (AAGATGATGTCCGGGTTTCAAGG) and 3′ (AATGTACTGGGGCATAAG), used to generate *Specc1l*^ΔEx4^ are depicted in Figure 1A, while exact locations were reported previously (Goering et al., 2021b). Correctly targeted ES cell clones were identified by PCR and sequencing and also checked by karyotyping. The resulting chimeric males were crossed to C57BL/6J females, and germline transmission confirmed by genotyping of offspringThe *Specc1l^fl^*, *Wnt1-Cre2*, and *ROSA^mT/mG^* reporter mice were maintained on a mixed C57BL/6J and FVB/NJ genetic background.

**Figure 1.**
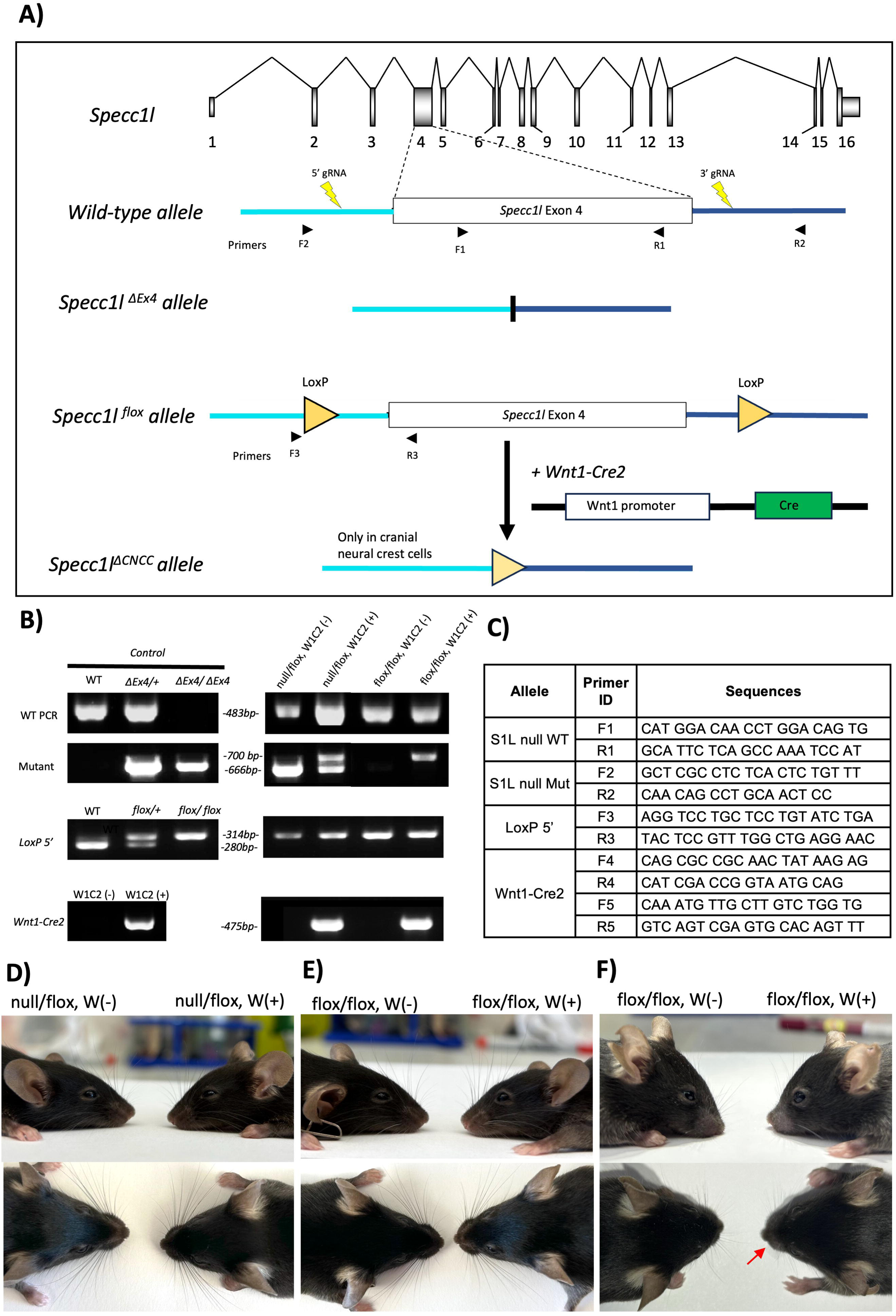
Generation of cranial neural crest specific *Specc1l* knockout. **(A)** Schematic representation of *Specc1l* locus, highlighting the exon 4 genomic region and the 5’ and 3’ guide RNAs (gRNAs). These gRNAs were previously used to generate the *Specc1l*^*ΔEx4*^ null allele. Here, these gRNAs were used to insert *loxP* sites flanking exon 4. The resulting *Specc1l* floxed allele was crossed with *Wnt1-Cre2* deleter strain to knockout *Specc1l* in cranial neural crest cells. Also shown are the approximate locations of the sequencing primers used for genotyping (B and C). **(B)** Genotyping analysis of the three alleles. Wild-type and mutant alleles were genotyped separately for Δ*Ex4*. **(C)** Sequence for primer pairs used for genotyping shown in B. Primer locations are also shown in the schematic of exon 4 in A. **(D-F)** Gross morphology of *Specc1l*^*ΔCNCC*^ mice (W+) compared to control littermates (W-). Both null/flox (D) and flox/flox (E) 10-week-old mice are shown with shortened frontonasal region. Also shown is an example of an 8-week-old flox/flox mutant (F) with a bent snout (arrow).

Conditional neural crest–specific deletion of *Specc1l* was achieved by crossing *Specc1l^fl^* mice with *Wnt1-Cre2* (RRID:IMSR_JAX:022501). The Wnt1-Cre2 allele was maintained on females, as *Specc1l^fl/+^;Wnt1-Cre2^+^,* and crossed with *Specc1l^fl/fl^*or *Specc1l^fl/null^* males to prevent male germline transmission that has been previously reported (Dinsmore et al., 2022). For lineage tracing, *ROSA^mT/mG^* (RRID:IMSR_JAX:007676) transgenic mice were used. Mice were housed in a pathogen-free facility, and all experimental procedures were conducted in accordance with protocols approved by the University of Kansas Medical Center Institutional Animal Care and Use Committee (IACUC).

### Genotyping

Tail biopsies were collected at weaning, and yolk sacs were obtained at the time of embryo harvesting. Genomic DNA was extracted using DirectPCR Lysis Reagent for Mouse Tail (Viagen, 102-T) or DirectPCR Lysis Reagent for Yolk Sac (Viagen, 202-Y), following the manufacturer’s instructions. PCR was performed using EconoTaq PLUS Master Mix to genotype *Specc1l* floxed, null, and *Cre* alleles, as well as *ROSA^mT/mG^* reporter configurations (**Fig.1B**), with primers listed in Figure 1C. PCR products were separated on a 1.5% agarose gel containing ethidium bromide and visualized under UV transillumination using a ChemiDoc imaging system (Bio-Rad Laboratories).

### Micro-computed tomography (microCT) visualization

Adult mice (7 −9 weeks in age) were euthanized in accordance with Kansas University Medical Center approved IACUC protocol (#23-11-35), fixed and stored in 4% PFA until imaging. Mice were imaged at 26-μm resolution using a Skyscan 1272 microCT scanner (Bruker). All images were acquired using the same settings (70 kV, 142 uA, 0.5 mm AI filter, 1000 ms exposure, 0.2° rotation step, 180° rotation, no frame averaging) for all specimens. Raw scan data were reconstructed using NRecon software (Bruker) and 3D rendered and segmented in Dragonfly (Comet Technologies Canada Inc.) and python using scikit-image and napari(van der Walt et al., 2014). Reconstruction, rendering and thresholding settings for segmentation were kept consistent between specimens. In cases where the standard threshold setting left some bones fused through tiny bridges: we masked the extra bone by using a higher threshold separated the two objects and then expanded the extra bone by three pixels. Dragonfly was used to quantify and visualize thickness, volume and density on segmented bones. Quantitative measurements were compared us54ing Student’s t test with Welch’s correction.

### Histology and immunofluorescence

Timed matings were established overnight and checked for vaginal plugs the following morning. Noon on the day a plug was detected was designated as embryonic day 0.5 (E0.5). Pregnant females were euthanized at the specified embryonic stages using IACUC-approved methods. Embryos at E9.5 and E13.5 were collected and fixed overnight in 4% paraformaldehyde (PFA) at 4°C. Samples were then cryoprotected sequentially in 15% and 30% sucrose solutions, each overnight at 4°C, and subsequently embedded in Optimal Cutting Temperature (OCT) compound for storage at −80°C. Prior to sectioning, samples were equilibrated at −20°C for several hours. Frozen tissues were sectioned on a cryostat at 10 μm thickness and mounted onto glass slides. Sections were allowed to equilibrate to room temperature (RT) for at least 30 minutes and kept in PBS to prevent drying.

For immunofluorescence, sections underwent antigen retrieval in preheated sodium citrate buffer for 20 minutes, followed by permeabilization in 0.5% Triton X-100 in PBS for 30 minutes. Sodium citrate buffer was made by dissolving 2.94g sodium citrate (Sigma, #S4641) in 1000 mL, adjust pH to 6.0 with 1N HCl then add 0.5 mL of Tween-20 (Fisher, #BP337). Blocking was performed at RT for 1 hour. Sections were then incubated overnight at 4°C with primary antibodies against SPECC1L N-terminus (1:250, Proteintech, 25390-1-AP), ARL13B (1:300, Proteintech, 17711-1-AP), Ki-67 (1:500, Cell Signaling, 12202), GLI1 (1:100, Cell Signaling, 2553S), GLI3 (1:100, R&D System, AF3690), Non-phospho (Active) β-catenin (1:500, Cell Signaling, 8814S), SOX10 (1:50, Proteintech, 10422-1-AP), SOX10 (1:30, Santa Cruz, sc-365692), β-catenin (1:250, Proteintech, 2677S), E-cadherin (1:400, Cell Signaling, 14472S). Secondary antibodies and stains were incubated for an hour: Goat anti-rabbit IgG (H+L) Alexa 647 (1:500, Invitrogen, A21245), Donkey anti-goat IgG (H+L) Alexa 647 (1:500, Invitrogen, A21447), Goat anti-rabbit IgG (H+L) Alexa 488 (1:500, Invitrogen, A11008), Goat anti-mouse IgG1 Alexa 488 (1:500, Invitrogen, A21121) and Acti-stain 670 phalloidin (1:200, Cytoskeleton, PHDN1-A). Note that for visualization of nuclear β-catenin, permeabilization was extended to 1 hour.

### Cilia length measurement

Cilia measurements were performed using images acquired primarily on a Nikon Eclipse Ti-E microscope equipped with an A1R confocal system. Z-stacks were captured with slices taken every 0.2 μm ensuring the full depth of each tissue section was captured. Maximum intensity projections (MIPs) were created for measurement. Cilia lengths were measured using ImageJ software and a segmented line tool was used to trace the cilium, following the method described by Jack & Avasthi(Jack et al., 2019). Measurements in pixels were converted to microns using the appropriate pixel-to-micron conversion factor for the objective used. Cilia length data were analyzed and graphed using GraphPad Prism, with mean ± 95% confidence intervals represented.

### Fluorescence Quantification

Cell fluorescence intensity was quantified using ImageJ/Fiji software. The corrected total cell fluorescence (CTCF) was calculated using the formula (Ansari et al., 2013):

CTCF=Integrated Density - (Area of selected area X Mean fluorescence of background readings).

For measurements, “Area”, “Integrated Density”, and “Mean Grey value” were selected under *Set Measurements*. Three background regions were measured to obtain an average background intensity. The region on interest (ROI) was then measured to obtain the integrated density value, and CTCF values were calculated accordingly. Final values were graphed using GraphPad Prism, with data represented as the mean ± standard deviation (SD).

## Results

### Frontonasal dysplasia upon loss of *Specc1l* in cranial neural crest cells

We previously reported a *Specc1l* null allele where we used CRISPR-Cas9 technology involving two guide RNAs (gRNAs) to delete exon 4 (**Fig.1A, *Specc1l***^Δ***Ex4***^ **or *Specc1l^null^***)(Goering et al., 2021b). To generate a conditional allele, we used the same gRNAs to insert *loxP* elements flanking exon 4 of *Specc1l* (**Fig.1A, *Specc1l^fl^***). The genotyping strategy for the null and floxed alleles (**Fig.1B**), as well as primer sequences used (**Fig.1C**) are described in more detail in the materials and methods. The *Wnt1-Cre2* allele was maintained on females to avoid known aberrant expression in the male germline, which can lead to unintended recombination in non-neural crest cells (Dinsmore et al.). To assess the role of *Specc1l* in CNCCs, *Specc1l^fl/+^; Wnt1-Cre2* females were crossed with *Specc1l^null/fl^* or *Specc1l^fl/fl^*males. Both *Specc1l^null/fl^;Wnt1-Cre2* and *Specc1l^fl/fl^;Wnt1-Cre2* progeny were assessed and are annotated in most figures. Since both mutant genotypes showed similar phenotypes, they are collectively referred to as *Specc1l*^*ΔCNCC*^ mice in the results. All *Specc1l*^*ΔCNCC*^ mutants exhibited a broad nasal bridge and a short snout (**Fig.1D,E**), 25% of which were asymmetrically leftward-bent (**Fig.1F, arrow**). Two *Specc1l^null/fl^;Wnt1-Cre2* embryos (∼3%) were observed to have a cleft palate. Together, these phenotypes suggest that loss of *Specc1l* in CNCCs can account for most of the craniofacial malformation associated with *SPECC1L-*related hypertelorism syndrome (Saadi et al., 2023).

### MicroCT analysis revealed frontal bone reduction and parietal bone increase in *Specc1l*^*ΔCNCC*^ mice

To obtain a better understanding of structural changes in *Specc1l*^*ΔCNCC*^ crania, we performed microCT analysis of 6–8-week-old mice (**Fig.2**). Both *Specc1l^null/fl^;Wnt1-Cre2* and *Specc1l^fl/fl^;Wnt1-Cre2* mice, with or without bent nose, are shown (**Fig.2A, Suppl. Fig.1**). Overall, we observed a significant decrease in frontal bone length (**Fig.2B,C; distance BC**), and a concomitant increase in parietal bone length (**Fig.2B,C; distance CD**). In contrast, skull bone width remained mostly similar, except a small increase at the position of the lambdoid suture (**Fig.2B,D; distance LM**). This small increase was sufficient to cause a significant increase in the ratio of width to total length measurement between wildtype and *Specc1l*^*ΔCNCC*^ mutant crania (**Fig.1E**), which is consistent with the brachycephaly associated with *SPECC1L-*related hypertelorism syndrome.

**Figure 2.**
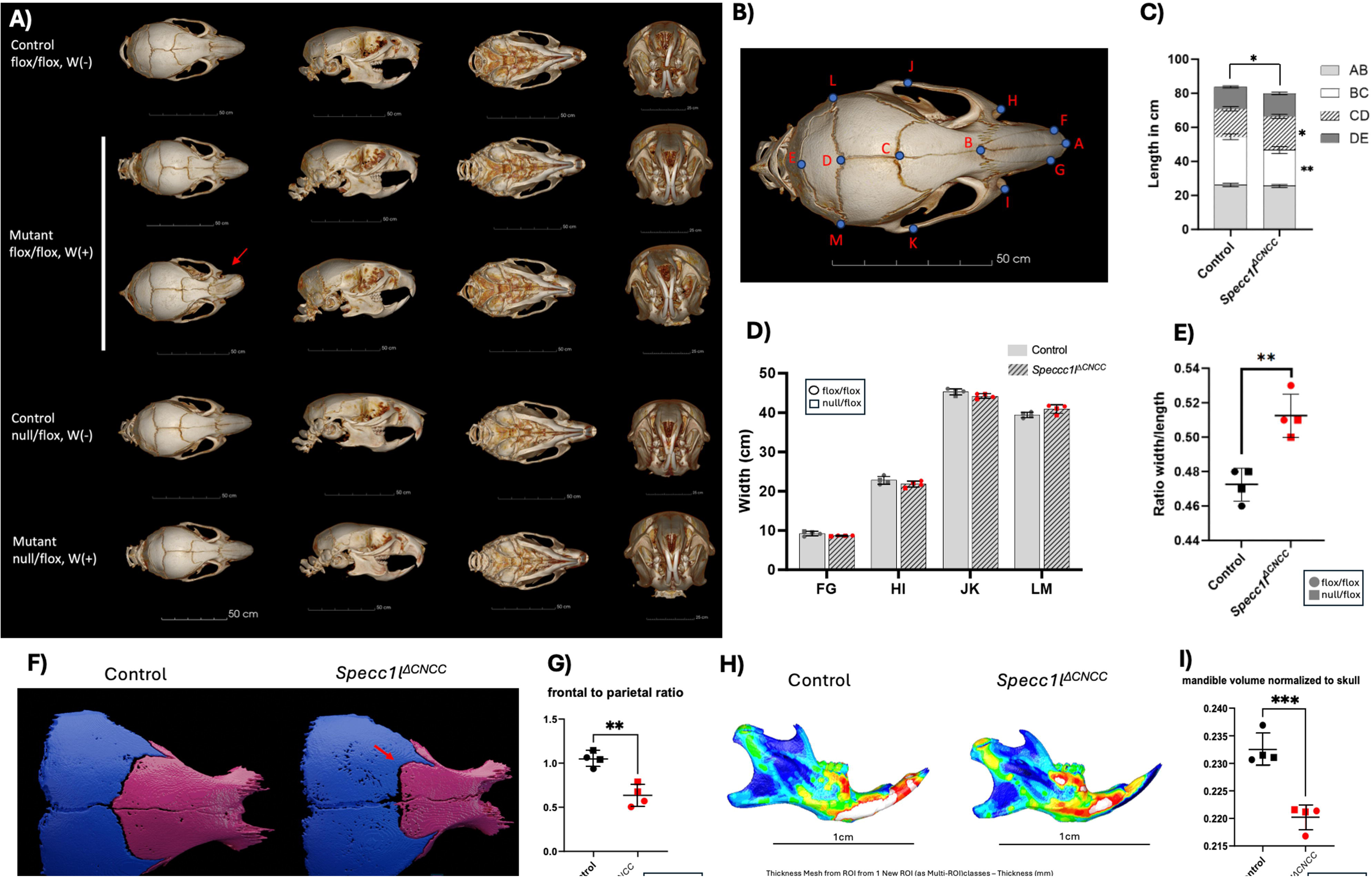
Craniofacial phenotypes of *Specc1l*^Δ*CNCC*^ mice. **(A)** Three-dimensional micro–computed tomography (microCT) reconstructions of postnatal mouse skulls shown in four standard orientations: dorsal, lateral, ventral, and anterior (left to right). **(B)** Anatomical landmarks used for craniofacial morphometric analyses in (C) and (D). **(C)** Quantification of skull length (cm) as a composite of nasal (AB), frontal (BC), parietal (CD) and interparietal (DE) bones. The overall skull length (AE) was shorter in the mutant mice (p<0.026), mainly driven by reduction in frontal bone (BC) length (p<0. 0015). In contrast, the parietal bone (CD) was longer in the mutants (p<0.0108). **(D)** Quantification of average skull width (cm) at the level of nasal tip (FG), nasal suture (HI), coronal suture (JK) and lambdoid suture (LM) did not show significant differences. **(E)** Ratio of skull width at lambdoid suture (LM) to length (AE) was significantly increased in the mutant mice (p<0.0028). **(F)** Segmentation of frontal and parietal bones showed a drastic change in coronal suture shape, which was more “box-like” in the mutant samples (arrow). **(G)** The differences in frontal and parietal bone sizes in the mutant skulls resulted in a significantly decreased frontal to parietal (p<0.0023). **(H)** Mandibular bone thickness maps did not show significant differences. **(I)** Normalized mandibular volume relative to total skull volume was significantly decreased in the mutant samples (p<0.0007). Data represent mean ± sd. Statistical significance was assessed using an unpaired two-tailed *t*-test.

We next examined the sizes of the cranial bones. There was a marked decrease in frontal bone area (**Fig.1F, magenta**). In contrast, parietal bone size was increased (**Fig.1F, blue**), resulting in a significant skewing of the frontal to parietal bone ratio (**Fig.1G**). In addition, the coronal suture shape appeared flatter and more ‘box-like’ in the mutant samples (**Fig.1F, arrow**). We also assessed regional bone volume and thickness (**Suppl. Fig.2**). We observed a significant reduction in mandible volume when normalized to the skull (**Fig.2H, I**). Malocclusion or incisor defects, however, were not observed (**Figs.2,3; Suppl. Figs.1,2**).

### *Specc1l*^*ΔCNCC*^ mice showed increased F-actin and reduced cell proliferation in the cranial mesenchyme

To assess the molecular underpinnings of the frontonasal dysplasia, we analyzed the cranial mesenchyme at the level of the developing frontal bone in E13.5 eombrtos (**Fig.3**). We also crossed the mutant alleles with *ROSA-mTmG* allele, which marks the Cre lineage traced CNCCs in green (**Suppl. Fig.3**). We confirmed that SPECC1L expression was diminished in the *Specc1l*^*ΔCNCC*^ cranial mesenchyme (**Fig.3A-C**). The *ROSA-mTmG* based CNCC-lineage was mapped only in the *Specc1l*^*ΔCNCC*^ mutant mice (**Fig.3**). Measurements were taken in the *Wnt1-Cre2* positive green region in the mutants, and in a comparable cranial mesenchyme region in controls (**Fig.3**). We next looked at levels of F-actin and cell proliferation via phalloidin and Ki-67 staining, respectively. SPECC1L has been shown to facilitate F-actin turnover (Saadi et al., 2011; Wilson et al., 2016; Hall et al., 2020; Goering et al., 2021b). Consistently, we observed an increase in F-actin staining in the *Specc1l*^*ΔCNCC*^ mutant cranial mesenchyme (**Fig.3D-F**). The cranial mesenchyme region in the mutant mice appeared narrower than in controls (**Fig.3A,D**). Thus, we examined cell proliferation and found it to be markedly decreased in the *Specc1l*^*ΔCNCC*^ mutant cranial mesenchyme (**Fig.3G-I**).

**Figure 3.**
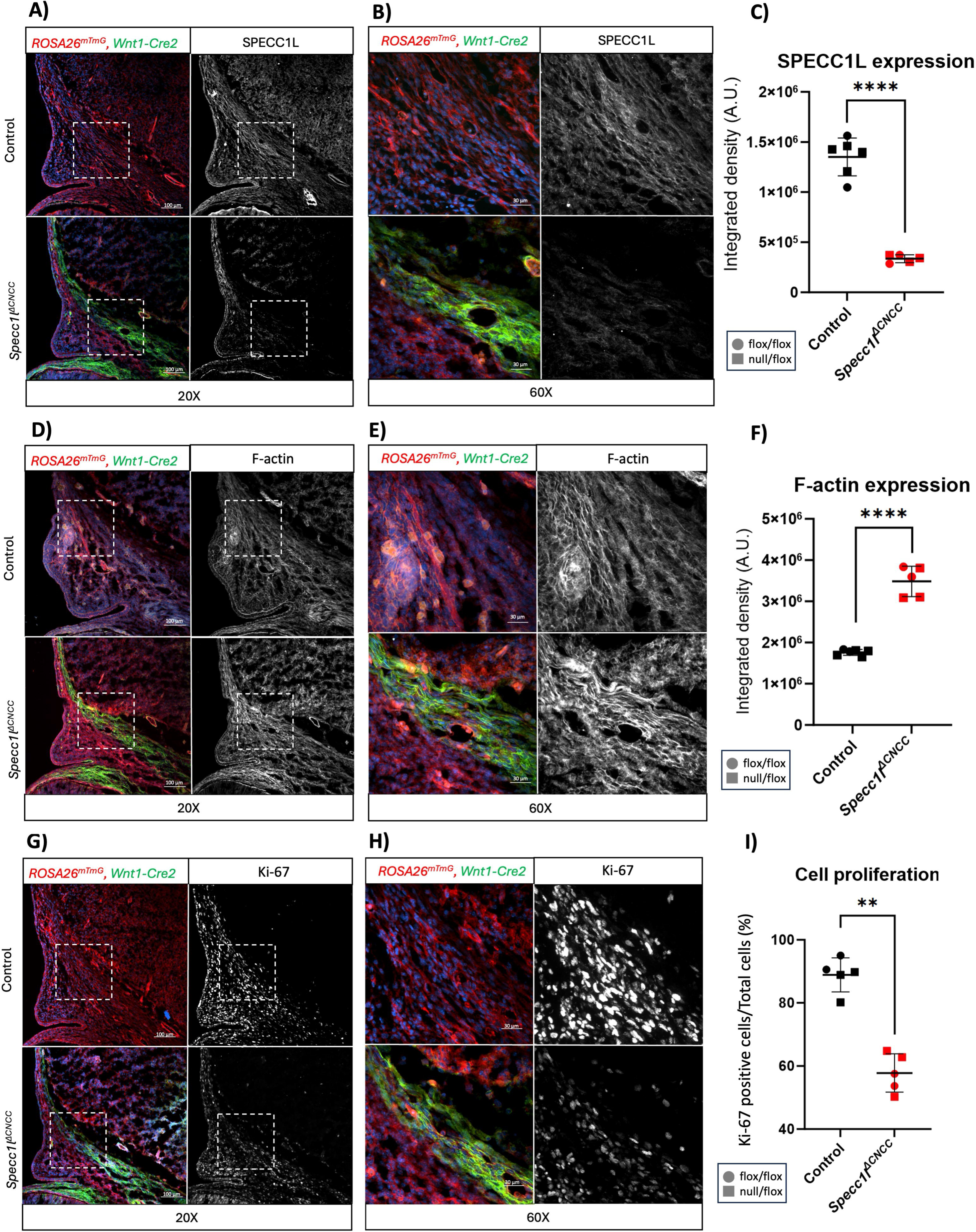
*Specc1l*^Δ*CNCC*^ cranial mesenchyme showed increased filamentous actin and decreased cell proliferation at E13.5. Immunofluorescence analysis of coronal sections at the level of frontal bone at E13.5. **(A-C)** Immunostaining for SPECC1L at 20x (A) and 60x (B) magnification of boxed region in A showed the expected loss of expression in cranial neural crest cell (CNCC) lineage (green) in *Specc1l*^*ΔCNCC*^ tissue. Control sample was *Wnt1-Cre2* negative. The corresponding quantification of integrated fluorescence intensity difference is shown (p<9.24E^-07^). **(D-F)** Phalloidin staining for filamentous actin (F-actin) at 20x (D) and 60x (E) magnification showed an increase in mutant CNCCs (F, p<1.24E^-06^). **(G-I)** Cell proliferation was assessed using Ki-67 immunolabeling. Images at 20X (G) and 60X (H) magnification, and quantitation of percent Ki-67 positive cells showed a decrease in cell proliferation in mutant CNCCs (I, p<0.0033). Data represent mean ± SD. Statistical significance was assessed using an unpaired two-tailed *t*-test, n=5.

### Altered ciliogenesis and hedgehog signaling in *Specc1l*^*ΔCNCC*^ cranial mesenchyme

We have shown that increased F-actin upon SPECC1L deficiency results in shortened cilia and increased Hh signaling in the palate at E13.5 (Hufft-Martinez et al., 2025). Thus, we expected cilia length and Hh signaling in the cranial mesenchyme to be perturbed. Indeed, cilia lengths were significantly decreased in *Specc1l*^*ΔCNCC*^ cranial mesenchyme at E13.5 (**Fig.4A,B**). Consistent with the ciliary defect, expression of GLI1, a downstream activator of hedgehog signaling, was increased in the *Specc1l*^*ΔCNCC*^ cranial mesenchyme (**Fig.4C,D**). Altered Hh signaling in ciliary mutants is known to affect canonical WNT signaling (Kurosaka et al., 2014). To this end we assessed expression of functionally active β-catenin, which was significantly decreased in *Specc1l*^*ΔCNCC*^ cranial mesenchyme (**Fig.4E,F**), consistent with the observed decrease in cell proliferation.

**Figure 4.**
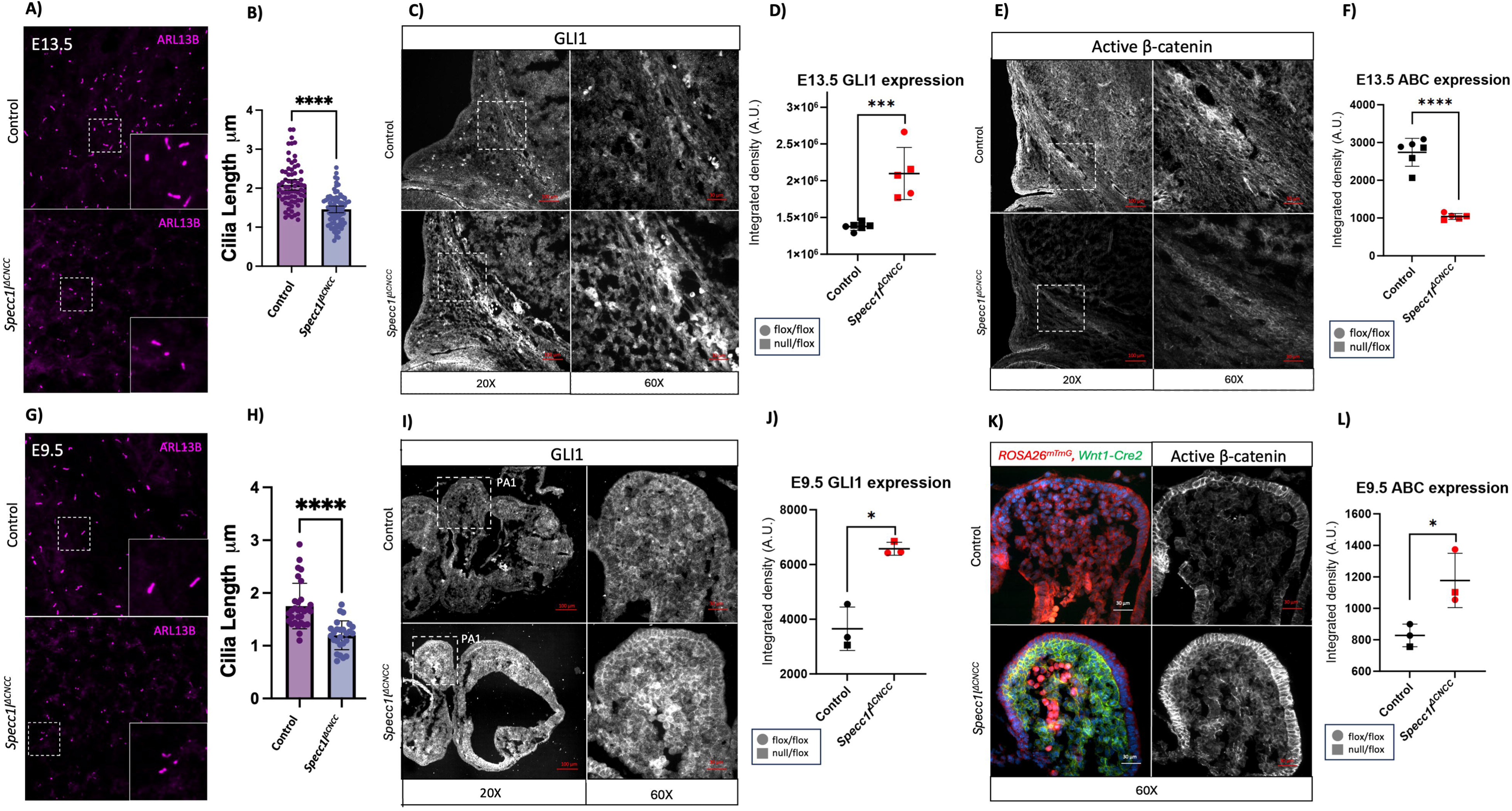
Shortened primary cilia and elevated hedgehog signaling in *Specc1l*^Δ*CNCC*^ cranial mesenchyme. Cranial mesenchyme was assessed in E13.5 coronal sections at the level of the frontal bone to assess cilia length using the ciliary membrane marker ARL13B **(A,B)**, hedgehog signaling using the downstream effector GLI1 immunostaining **(C,D)**, and canonical WNT signaling using active β-catenin immunostaining **(E,F)**. Boxed regions are magnified in insets (A) or in panels to the right (C,E). Cilia length measurement (B) showed a significant reduction in E13.5 *Specc1l*^*ΔCNCC*^ cranial mesenchymal cells (n=85), compared with control cells (n=70). GLI1 levels were increased in the mutant samples (D), while active β-catenin levels were decreased (F). To determine if these changes were potentially causal, E9.5 sections through the first pharyngeal arch (PA1) were assessed. The cilia length was decreased in comparison between control (n=20) and *Specc1l*^*ΔCNCC*^ (n=53) **(G,H)**, and GLI1 levels increased **(I,J)** in *Specc1l*^*ΔCNCC*^ mutant mesenchyme, similarly to E13.5 cranial mesenchyme. However, active β-catenin levels were increased in the E9.5 mutant mesenchyme **(K,L)**, in contrast to E13.5 cranial mesenchyme. Cilia length analysis represents mean ± 95% CI. Remaining analyses represent mean ± SD. Statistical significance was assessed using an unpaired two-tailed *t*-test; n=5 for **D**, **F** and n=3 for **J**, **M** (* p<0.05, ** p<0.01, *** p<0.001, **** p<0.0001).

We next asked whether these changes were potentially causal. We evaluated the *Specc1l*^*ΔCNCC*^ mutant embryos at E9.5. We observed a similar shortening of cilia (**Fig.4G,H**) and increased GLI1 expression (**Fig.4I,J**) in E9.5 facial prominences of *Specc1l*^*ΔCNCC*^ mutant embryos. However, we observed increased staining of active β-catenin in the mutant tissue (**Fig.4K,L**), which correlated with increased Ki-67 staining (**Suppl. Fig.4**). Our data suggest that shortened cilia and increased hedgehog signaling are early events upon *Specc1l* loss. β-catenin function, in contrast, is differentially affected over cranial mesenchyme development, likely due to the ciliary signaling defect. We previously reported ectopically stabilized cell-cell adhesions in migratory CNCCs in a globally *Specc1l*-deficient allele (Bertol et al., 2022). We found similarly increased expression of adherens junction markers, E-cadherin and β-catenin, SOX10-positive migratory CNCCs in *Specc1l*^*ΔCNCC*^ mutant embryos at E9.5 (**Suppl. Fig.5**). Together, these findings support the conclusion that *Specc1l* deficiency disrupts cilia-based signaling early in CNCC function affecting migration, signaling, and differentiation.

## Discussion

Frontonasal dysplasia is a collection of disorders with variable effects (Sedano et al., 1970; Sedano and Gorlin, 1988; Farlie et al., 2016). *SPECC1L-*related syndrome is also referred to as TBHS1 (OMIM: 145420) or brachycephalofrontonasal dysplasia (ORPHA:1519), which involves shortening of the cranium with posterior widening. Our *Specc1l*^*ΔCNCC*^ mutant mice showed a similar phenotype of cranial shortening due to reduction in frontal bone size, and widening at the lambdoid suture, likely due to abnormal compensatory growth of the parietal bone. Nasal bone architecture was also altered, and ∼3% of the mutant mice developed cleft palate. Thus, the major craniofacial features of the *SPECC1L-*related hypertelorism syndrome appear to be CNCC-derived. While we did observe an increase in hedgehog signaling, which regulates midfacial growth, we did not observe an increase in the inter-canthal distance in the *Specc1l*^*ΔCNCC*^ mutant mice. Thus, the most canonical hypertelorism feature of the *SPECC1L-*related syndrome likely involves function of SPECC1L in cells beyond CNCCs.

A striking feature of *Specc1l*^*ΔCNCC*^ mutant crania was the change in the coronal suture shape, which appeared more ‘box-like’ with a sharp transition between frontal and parietal bones (**Fig.1F**). Similarly altered coronal sutures have been observed in *Twist1^+/-^*mice (Bialek et al., 2004; Bertol et al., 2022). Specifically, Teng *et al*. (Teng et al., 2018) showed that *Twist1* haploinsufficiency in the mesoderm (*Twi1^fl/+^;Mesp1-Cre*) leads to the exact same change in the coronal suture shape. They also showed an increase in parietal bone size and a concomitant decrease in frontal bone size in *Twi1^fl/+^;Mesp1-Cre* mice, exactly similar to our *Specc1l*^*ΔCNCC*^ mutant mice. However, when Teng *et al*. deleted a copy of *Twist1* in the neural crest, the *Twi1^fl/+^;Wnt1-Cre* mice exhibited an increase in frontal bone and a decrease in parietal bone size. *TWIST1* and *SPECC1L* functions intersect in at least two aspects. *TWIST1* mutations are associated with syndromes primarily characterized by craniosynostosis and variably by facial dysmorphism, including cleft palate (Topa et al., 2020; Bertol et al., 2022). Similarly, three patients with *SPECC1L-*related hypertelorism syndrome also manifested craniosynostosis (Bhoj et al., 2019). Additionally, we previously reported that TWIST1 can bind directly to *Specc1l* putative intronic regulatory elements, and that *Specc1l* expression was decreased in early embryonic tissue from *Twist1* mutants (Bertol et al., 2022). These observations suggest a complementary relationship between *Specc1l* and *Twist1*.

In addition to *Twist1,* combinatorial reduction in *Msx1* and *Msx2* dosage in the CNCCs affected frontal bone formation (Roybal et al., 2010). Heterozygous loss of *Efnb1* in CNCCs alone, or in combination with *Efnb2* heterozygosity, also affected frontal bone development (Davy et al., 2006). Loss of *Fgfr1* in the CNCCs did not appear to change the frontal bone size but led to heterotopic osteogenesis (Kawai et al., 2019). Both ephrin and FGF signaling also affect cilia, and MSX1/2 function downstream of Hh signaling in the calvarial bone (Kunova Bosakova et al., 2019; Cho et al., 2025; Loukil et al., 2025). In contrast, loss of *Mid1* in CNCCs resulted in an increase in both frontal and nasal bones (Liang et al., 2023). *MID1* mutations result in X-linked Opitz GBBB syndrome (OMIM:300000) with a phenotypic spectrum similar to that of *SPECC1L*-related syndrome, including hypertelorism, cleft lip/palate, cardiac defects and hypospadias(Opitz, 1987; So et al., 2005). In fact, *SPECC1L* mutations have been identified in patients characterized by non-X-linked Opitz GBBB syndrome (Kruszka et al., 2015).

The compensatory changes in frontal and parietal bone sizes have also been reported in mouse mutants in hedgehog signaling pathway. In the *Fuz* mutant mice, the frontal bone expands at the expense of the parietal bone, which could be rescued with reduction in *Fgf8* levels (Tabler et al., 2016). FUZ is an essential regulator of ciliogenesis, where it controls the processing of GLI3 full length (GLI3FL) into its cleaved repressor form (GLI3R). In *Fuz* mutants, there is an increase in GLI3FL while GLI1 levels either remain unchanged or decrease depending on the tissue. In our *Specc1l*^*ΔCNCC*^ mutant tissue, GLI3 levels are decreased upon immunostaining (**Suppl. Fig.6**), however, we could not distinguish between GLI3FL and GLI3R levels. While shortened cilia and increased GLI1 levels were observed at both E9.5 and E13.5 in our *Specc1l*^*ΔCNCC*^ mutant cranial mesenchyme, active β-catenin levels differed (**Fig.4**). Hedgehog signaling normally promotes cell proliferation. However, Mak *et al*. (2008) reported that increased hedgehog signaling in mature osteoblasts resulted in ectopically induced osteoclast differentiation leading to bone loss (Mak et al., 2008). Thus, increased hedgehog signaling in *Specc1l*^*ΔCNCC*^ mutant cranial mesenchyme may initially promote proliferation but may eventually result in abnormal differentiation and frontonasal bone malformation. Our data suggest that loss of *Specc1l* in CNCCs results in a shift in the neural crest-mesoderm interface in a direction opposite to that of the *Fuz* mutant.

Overall, loss of SPECC1L results in increased F-actin in CNCCs, which results in shortened cilia and increased Hh signaling, affecting CNCC migration and differentiation. The ciliary and Hh defects also affect canonical WNT signaling and cell proliferation, resulting in imbalanced growth of frontal and parietal bones.

## Author Contributions

IS conceived the experiments. AJT, BMH, JPG and IS designed the experiments. AJT, BMH, LM, and JPG performed the experiments. LM, SM, SN and PT performed the microCT scans and analyses. AJT, BMH, LM, DNT and IS wrote the paper. JPG and PT edited the manuscript. All authors reviewed the manuscript.

## Acknowledgments

We want to thank Dr. Jay Vivian, the KUMC Transgenic Core facility, and Luke Wenger for their help in the design and generation of the *Specc1l* conditional allele. We would like to thank Stephanie Nowotarski and Melania McClain in the Stowers Electron and Light Microscopy Technology Center for assistance with the micro-CT instrument and data visualization.

## Data Availability Statement

The original contributions presented in the study are included in the article/Supplementary Material, further inquiries can be directed to the corresponding author. Original data underlying this manuscript that was generated at the Stowers Institute for Medical Research can be accessed from the Stowers Original Data Repository at https://www.stowers.org/research/publications.

## References

Achilleos, A., Huffman, N.T., Marcinkiewicyz, E., Seidah, N.G., Chen, Q., Dallas, S.L., et al. (2015). MBTPS1/SKI-1/S1P proprotein convertase is required for ECM signaling and axial elongation during somitogenesis and vertebral developmentdagger. Hum Mol Genet 24(10), 2884–2898. doi: 10.1093/hmg/ddv050.

Ahlgren, S.C., and Bronner-Fraser, M. (1999). Inhibition of sonic hedgehog signaling in vivo results in craniofacial neural crest cell death. Curr Biol 9(22), 1304–1314. doi: 10.1016/s0960-9822(00)80052-4.

Ansari, N., Müller, S., Stelzer, E.H., and Pampaloni, F. (2013). Quantitative 3D cell-based assay performed with cellular spheroids and fluorescence microscopy. Methods Cell Biol 113, 295–309. doi: 10.1016/b978-0-12-407239-8.00013-6.

Assis, L.H., Silva-Junior, R.M., Dolce, L.G., Alborghetti, M.R., Honorato, R.V., Nascimento, A.F., et al. (2017). The molecular motor Myosin Va interacts with the cilia-centrosomal protein RPGRIP1L. Sci Rep 7, 43692. doi: 10.1038/srep43692.

Bertol, J.W., Johnston, S., Ahmed, R., Xie, V.K., Hubka, K.M., Cruz, L., et al. (2022). TWIST1 interacts with β/δ-catenins during neural tube development and regulates fate transition in cranial neural crest cells. Development 149(15). doi: 10.1242/dev.200068.

Bhoj, E.J., Haye, D., Toutain, A., Bonneau, D., Nielsen, I.K., Lund, I.B., et al. (2019). Phenotypic spectrum associated with SPECC1L pathogenic variants: new families and critical review of the nosology of Teebi, Opitz GBBB, and Baraitser-Winter syndromes. Eur J Med Genet 62(12), 103588. doi: 10.1016/j.ejmg.2018.11.022.

Bhoj, E.J., Li, D., Harr, M.H., Tian, L., Wang, T., Zhao, Y., et al. (2015). Expanding the SPECC1L mutation phenotypic spectrum to include Teebi hypertelorism syndrome. Am J Med Genet A 167a(11), 2497–2502. doi: 10.1002/ajmg.a.37217.

Bialek, P., Kern, B., Yang, X., Schrock, M., Sosic, D., Hong, N., et al. (2004). A twist code determines the onset of osteoblast differentiation. Dev Cell 6(3), 423–435. doi: 10.1016/s1534-5807(04)00058-9.

Bisgrove, B.W., and Yost, H.J. (2006). The roles of cilia in developmental disorders and disease. Development 133(21), 4131–4143. doi: 10.1242/dev.02595.

Bleicher, P., Sciortino, A., and Bausch, A.R. (2020). The dynamics of actin network turnover is self-organized by a growth-depletion feedback. Sci Rep 10(1), 6215. doi: 10.1038/s41598-020-62942-8.

Briscoe, J., and Thérond, P.P. (2013). The mechanisms of Hedgehog signalling and its roles in development and disease. Nat Rev Mol Cell Biol 14(7), 416–429. doi: 10.1038/nrm3598.

Chang, C.F., Chang, Y.T., Millington, G., and Brugmann, S.A. (2016). Craniofacial Ciliopathies Reveal Specific Requirements for GLI Proteins during Development of the Facial Midline. PLoS Genet 12(11), e1006351. doi: 10.1371/journal.pgen.1006351.

Chiang, C., Litingtung, Y., Lee, E., Young, K.E., Corden, J.L., Westphal, H., et al. (1996). Cyclopia and defective axial patterning in mice lacking Sonic hedgehog gene function. Nature 383(6599), 407–413. doi: 10.1038/383407a0.

Cho, J.G., Hah, Y., Yun, E., Ka, H.I., Lee, A., Han, S., et al. (2025). Loss of primary cilia promotes EphA2-mediated endothelial-to-mesenchymal transition in the ovarian tumor microenvironment. Mol Oncol 19(10), 2951–2966. doi: 10.1002/1878-0261.70057.

Cordero, D.R., Brugmann, S., Chu, Y., Bajpai, R., Jame, M., and Helms, J.A. (2011). Cranial neural crest cells on the move: their roles in craniofacial development. Am J Med Genet A 155a(2), 270-279. doi: 10.1002/ajmg.a.33702.

Davy, A., Bush, J.O., and Soriano, P. (2006). Inhibition of gap junction communication at ectopic Eph/ephrin boundaries underlies craniofrontonasal syndrome. PLoS Biol 4(10), e315. doi: 10.1371/journal.pbio.0040315.

Dinsmore, C.J., Ke, C.Y., and Soriano, P. (2022). The Wnt1-Cre2 transgene is active in the male germline. Genesis 60(3), e23468. doi: 10.1002/dvg.23468.

Farlie, P.G., Baker, N.L., Yap, P., and Tan, T.Y. (2016). Frontonasal Dysplasia: Towards an Understanding of Molecular and Developmental Aetiology. Mol Syndromol 7(6), 312–321. doi: 10.1159/000450533.

Goering, J.P., Isai, D.G., Hall, E.G., Wilson, N.R., Kosa, E., Wenger, L.W., et al. (2021a). SPECC1L-deficient primary mouse embryonic palatal mesenchyme cells show speed and directionality defects. Sci Rep 11(1), 1452. doi: 10.1038/s41598-021-81123-9.

Goering, J.P., Wenger, L.W., Stetsiv, M., Moedritzer, M., Hall, E.G., Isai, D.G., et al. (2021b). In-frame deletion of SPECC1L microtubule association domain results in gain-of-function phenotypes affecting embryonic tissue movement and fusion events. Hum Mol Genet 31(1), 18–31. doi: 10.1093/hmg/ddab211.

Goetz, S.C., and Anderson, K.V. (2010). The primary cilium: a signalling centre during vertebrate development. Nat Rev Genet 11(5), 331–344. doi: 10.1038/nrg2774.

Hall, E.G., Wenger, L.W., Wilson, N.R., Undurty-Akella, S.S., Standley, J., Augustine-Akpan, E.A., et al. (2020). SPECC1L regulates palate development downstream of IRF6. Hum Mol Genet 29(5), 845–858. doi: 10.1093/hmg/ddaa002.

Han, J., Mayo, J., Xu, X., Li, J., Bringas, P., Jr., Maas, R.L., et al. (2009). Indirect modulation of Shh signaling by Dlx5 affects the oral-nasal patterning of palate and rescues cleft palate in Msx1-null mice. Development 136(24), 4225–4233. doi: 10.1242/dev.036723.

Hufft-Martinez, B.M., Thalman, D.N., Tran, A.J., Goering, J.P., Stetsiv, M., Moedritzer, M., et al. (2025). Genetic interaction of *Specc1l* and *Thm1* reveals cytoskeletal - ciliary crosstalk. bioRxiv, 2025.2011.2003.686369. doi: 10.1101/2025.11.03.686369.

Jack, B., Mueller, D.M., Fee, A.C., Tetlow, A.L., and Avasthi, P. (2019). Partially Redundant Actin Genes in Chlamydomonas Control Transition Zone Organization and Flagellum-Directed Traffic. Cell Rep 27(8), 2459–2467.e2453. doi: 10.1016/j.celrep.2019.04.087.

Jeong, J., Mao, J., Tenzen, T., Kottmann, A.H., and McMahon, A.P. (2004). Hedgehog signaling in the neural crest cells regulates the patterning and growth of facial primordia. Genes Dev 18(8), 937–951. doi: 10.1101/gad.1190304.

Kawai, M., Herrmann, D., Fuchs, A., Cheng, S., Ferrer-Vaquer, A., Götz, R., et al. (2019). Fgfr1 conditional-knockout in neural crest cells induces heterotopic chondrogenesis and osteogenesis in mouse frontal bones. Med Mol Morphol 52(3), 156–163. doi: 10.1007/s00795-018-0213-z.

Kruszka, P., Li, D., Harr, M.H., Wilson, N.R., Swarr, D., McCormick, E.M., et al. (2015). Mutations in SPECC1L, encoding sperm antigen with calponin homology and coiled-coil domains 1-like, are found in some cases of autosomal dominant Opitz G/BBB syndrome. J Med Genet 52(2), 104–110. doi: 10.1136/jmedgenet-2014-102677.

Kunova Bosakova, M., Nita, A., Gregor, T., Varecha, M., Gudernova, I., Fafilek, B., et al. (2019). Fibroblast growth factor receptor influences primary cilium length through an interaction with intestinal cell kinase. Proc Natl Acad Sci U S A 116(10), 4316-4325. doi: 10.1073/pnas.1800338116.

Kuriyama, S., and Mayor, R. (2008). Molecular analysis of neural crest migration. Philos Trans R Soc Lond B Biol Sci 363(1495), 1349–1362. doi: 10.1098/rstb.2007.2252.

Kurosaka, H., Iulianella, A., Williams, T., and Trainor, P.A. (2014). Disrupting hedgehog and WNT signaling interactions promotes cleft lip pathogenesis. J Clin Invest 124(4), 1660–1671. doi: 10.1172/jci72688.

Liang, C., Profico, A., Buzi, C., Khonsari, R.H., Johnson, D., O’Higgins, P., et al. (2023). Normal human craniofacial growth and development from 0 to 4 years. Sci Rep 13(1), 9641. doi: 10.1038/s41598-023-36646-8.

Loukil, A., Ebright, E., Hamdi, K., Menzel, E., Uezu, A., Gao, Y., et al. (2025). Identification of New Ciliary Signaling Pathways in the Brain and Insights into Neurological Disorders. J Neurosci 45(33). doi: 10.1523/jneurosci.0800-24.2025.

Mak, K.K., Bi, Y., Wan, C., Chuang, P.T., Clemens, T., Young, M., et al. (2008). Hedgehog signaling in mature osteoblasts regulates bone formation and resorption by controlling PTHrP and RANKL expression. Dev Cell 14(5), 674–688. doi: 10.1016/j.devcel.2008.02.003.

Mansouri, A., Stoykova, A., Torres, M., and Gruss, P. (1996). Dysgenesis of cephalic neural crest derivatives in Pax7-/- mutant mice. Development 122(3), 831–838. doi: 10.1242/dev.122.3.831.

Mehta, V., Decan, N., Ooi, S., Gaudreau-Lapierre, A., Copeland, J.W., and Trinkle-Mulcahy, L. (2023). SPECC1L binds the myosin phosphatase complex MYPT1/PP1β and can regulate its distribution between microtubules and filamentous actin. J Biol Chem 299(2), 102893. doi: 10.1016/j.jbc.2023.102893.

Opitz, J.M. (1987). G syndrome (hypertelorism with esophageal abnormality and hypospadias, or hypospadias-dysphagia, or “Opitz-Frias” or “Opitz-G” syndrome)--perspective in 1987 and bibliography. Am J Med Genet 28(2), 275–285. doi: 10.1002/ajmg.1320280203.

Parker, F., Baboolal, T.G., and Peckham, M. (2020). Actin Mutations and Their Role in Disease. Int J Mol Sci 21(9). doi: 10.3390/ijms21093371.

Pollard, T.D., and Cooper, J.A. (2009). Actin, a central player in cell shape and movement. Science 326(5957), 1208–1212. doi: 10.1126/science.1175862.

Roybal, P.G., Wu, N.L., Sun, J., Ting, M.C., Schafer, C.A., and Maxson, R.E. (2010). Inactivation of Msx1 and Msx2 in neural crest reveals an unexpected role in suppressing heterotopic bone formation in the head. Dev Biol 343(1-2), 28–39. doi: 10.1016/j.ydbio.2010.04.007.

Saadi, I., Alkuraya, F.S., Gisselbrecht, S.S., Goessling, W., Cavallesco, R., Turbe-Doan, A., et al. (2011). Deficiency of the cytoskeletal protein SPECC1L leads to oblique facial clefting. Am J Hum Genet 89(1), 44–55. doi: 10.1016/j.ajhg.2011.05.023.

Saadi, I., Goering, J.P., Hufft-Martinez, B.M., and Tran, P.V. (2023). SPECC1L: a cytoskeletal protein that regulates embryonic tissue dynamics. Biochem Soc Trans 51(3), 949–958. doi: 10.1042/BST20220461.

Sandell, L.L., Iulianella, A., Melton, K.R., Lynn, M., Walker, M., Inman, K.E., et al. (2011). A phenotype-driven ENU mutagenesis screen identifies novel alleles with functional roles in early mouse craniofacial development. Genesis 49(4), 342–359. doi: 10.1002/dvg.20727.

Sedano, H.O., Cohen, M.M., Jr., Jirasek, J., and Gorlin, R.J. (1970). Frontonasal dysplasia. J Pediatr 76(6), 906–913. doi: 10.1016/s0022-3476(70)80374-2.

Sedano, H.O., and Gorlin, R.J. (1988). Frontonasal malformation as a field defect and in syndromic associations. Oral Surg Oral Med Oral Pathol 65(6), 704–710. doi: 10.1016/0030-4220(88)90014-x.

So, J., Suckow, V., Kijas, Z., Kalscheuer, V., Moser, B., Winter, J., et al. (2005). Mild phenotypes in a series of patients with Opitz GBBB syndrome with MID1 mutations. Am J Med Genet A 132A(1), 1-7. doi: 10.1002/ajmg.a.30407.

Tabler, J.M., Rice, C.P., Liu, K.J., and Wallingford, J.B. (2016). A novel ciliopathic skull defect arising from excess neural crest. Dev Biol 417(1), 4-10. doi: 10.1016/j.ydbio.2016.07.001.

Teng, C.S., Ting, M.C., Farmer, D.T., Brockop, M., Maxson, R.E., and Crump, J.G. (2018). Altered bone growth dynamics prefigure craniosynostosis in a zebrafish model of Saethre-Chotzen syndrome. Elife 7. doi: 10.7554/eLife.37024.

Tobin, J.L., Di Franco, M., Eichers, E., May-Simera, H., Garcia, M., Yan, J., et al. (2008). Inhibition of neural crest migration underlies craniofacial dysmorphology and Hirschsprung’s disease in Bardet-Biedl syndrome. Proc Natl Acad Sci U S A 105(18), 6714–6719. doi: 10.1073/pnas.0707057105.

Topa, A., Rohlin, A., Andersson, M.K., Fehr, A., Lovmar, L., Stenman, G., et al. (2020). NGS targeted screening of 100 Scandinavian patients with coronal synostosis. Am J Med Genet A 182(2), 348-356. doi: 10.1002/ajmg.a.61427.

Trainor, P.A. (2005). Specification and patterning of neural crest cells during craniofacial development. Brain Behav Evol 66(4), 266-280. doi: 10.1159/000088130.

van der Walt, S., Schönberger, J.L., Nunez-Iglesias, J., Boulogne, F., Warner, J.D., Yager, N., et al. (2014). scikit-image: image processing in Python. PeerJ 2, e453. doi: 10.7717/peerj.453.

Wilson, N.R., Olm-Shipman, A.J., Acevedo, D.S., Palaniyandi, K., Hall, E.G., Kosa, E., et al. (2016). SPECC1L deficiency results in increased adherens junction stability and reduced cranial neural crest cell delamination. Sci Rep 6, 17735. doi: 10.1038/srep17735.

Xu, J., Iyyanar, P.P.R., Lan, Y., and Jiang, R. (2023). Sonic hedgehog signaling in craniofacial development. Differentiation 133, 60–76. doi: 10.1016/j.diff.2023.07.002.

