## Supplemental Figures for "Loss of SPECC1L in cranial neural crest cells results in increased hedgehog signaling and frontonasal dysplasia"

### *Supplementary Material*

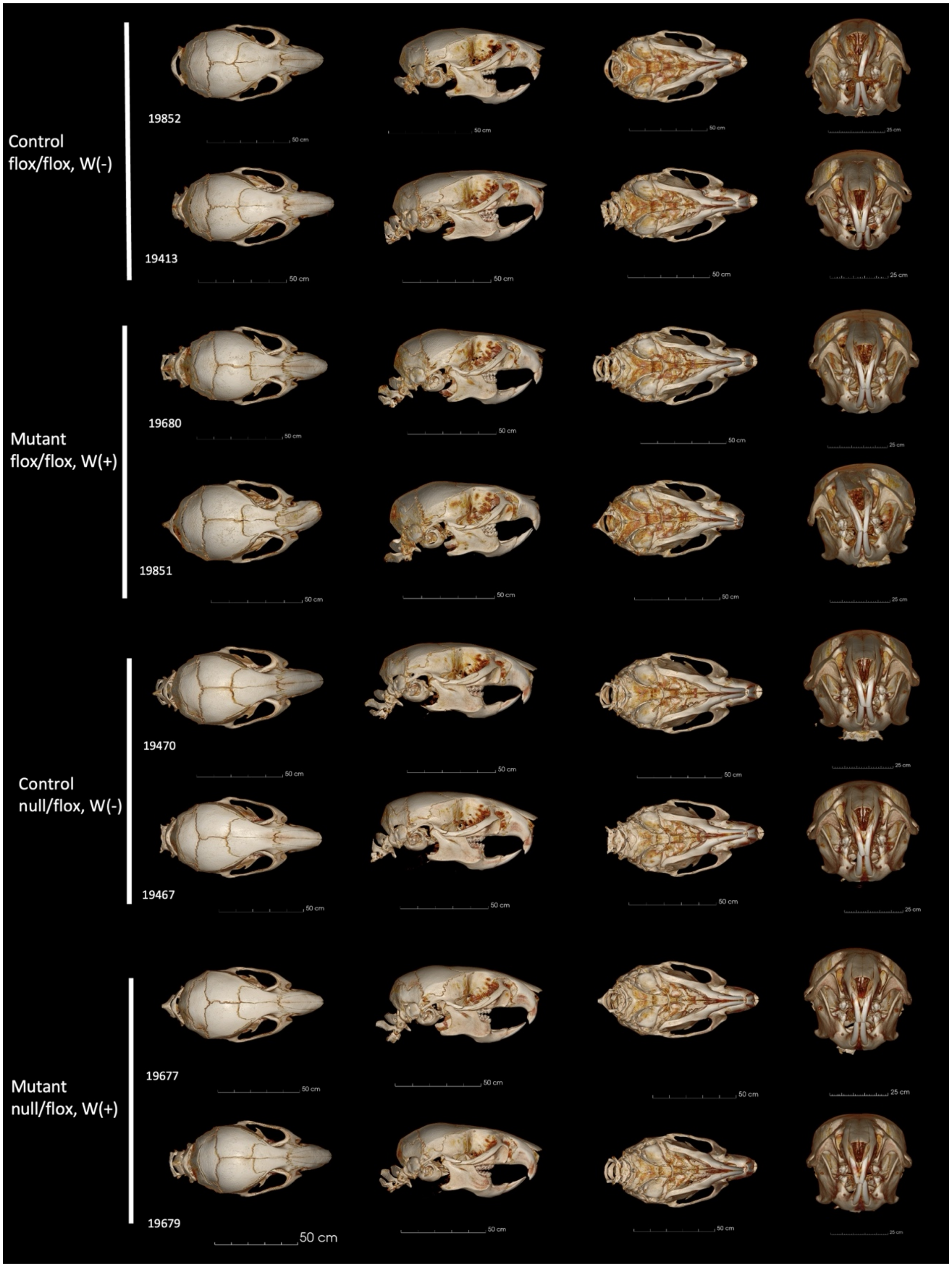


**Supplementary Figure 1**: Three-dimensional micro–computed tomography (microCT) reconstructions of all samples are presented in four orientations: dorsal, lateral, ventral, and anterior (from left to right).

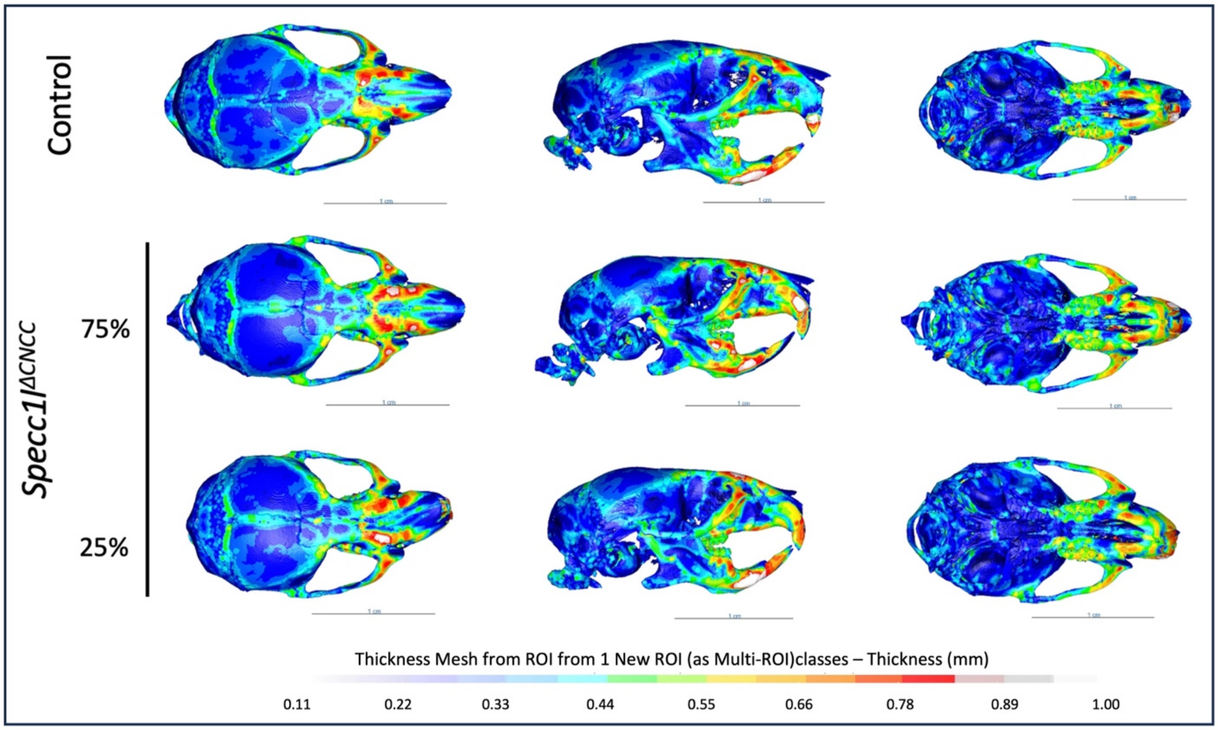


**Supplementary Figure 2:** Bone thickness heatmaps reveal regional heterogeneity in mutant skulls compared to control in dorsal, lateral and ventral orientations (left to right). *Specc1l*$^{\Delta CNCC}$ mice without and with bent snout are shown.

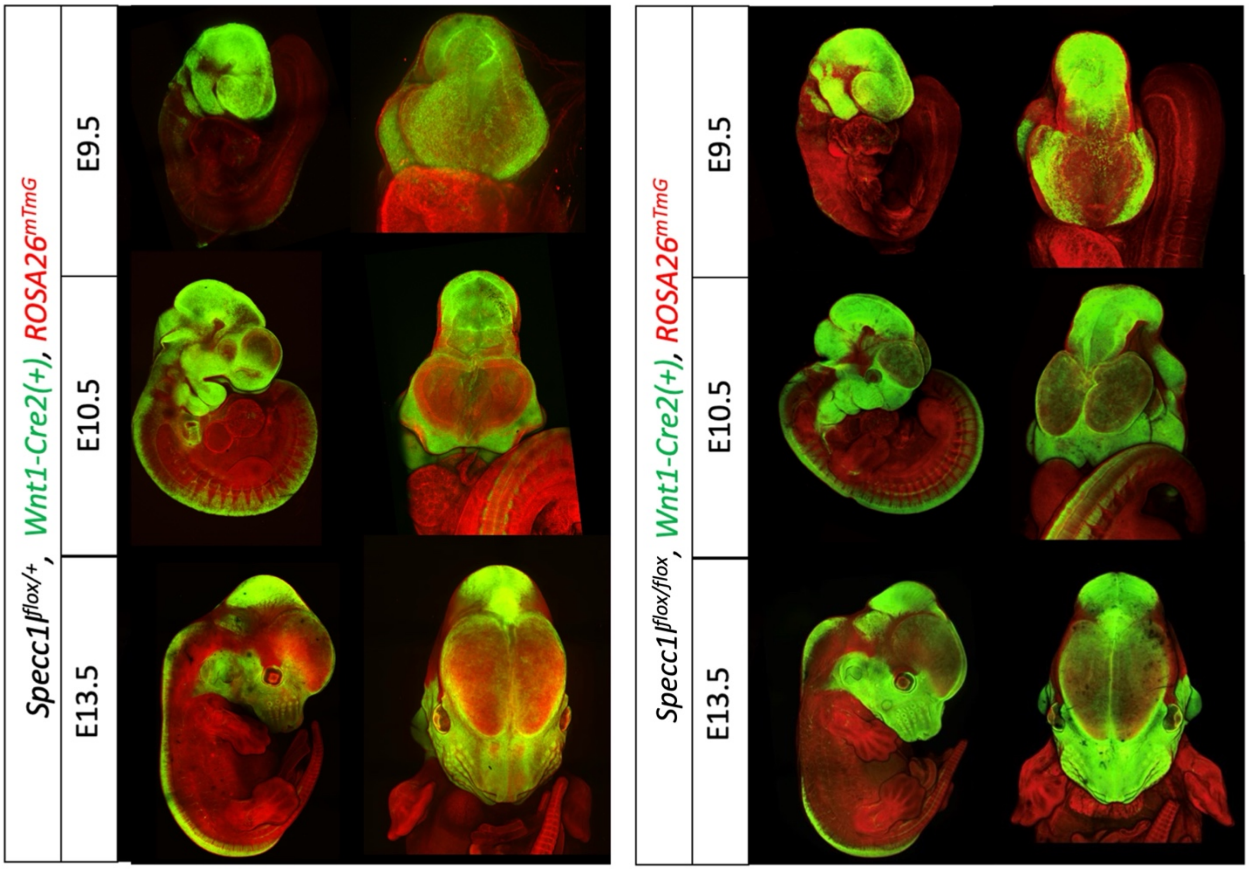


**Supplementary Figure 3:** To visualize neural crest cell distribution and confirm the spatial pattern of *Specc1l* deletion during craniofacial morphogenesis, *Wnt1-Cre2*;*ROSA*$^{mT/mG}$ reporter mice were examined at multiple developmental stages.

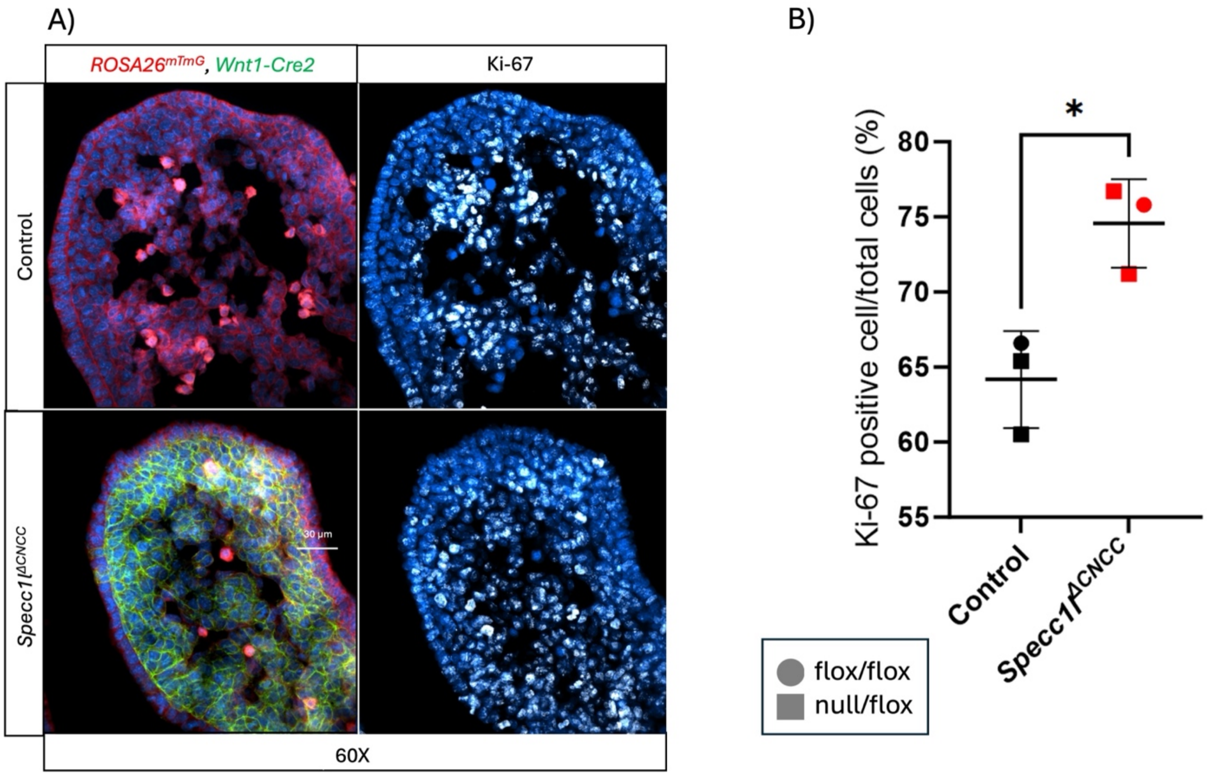


**Supplementary Figure 4:** Cell proliferation was assessed in E9.5 first pharyngeal arch using Ki-67 immunolabeling **(A)**. Percentage of Ki-67 positive cells over total nuclei showed a significant increase in *Specc1l*$^{\Delta CNCC}$ mesenchyme **(B)**. Data represent mean ± SD. Statistical significance was assessed using an unpaired two-tailed *t*-test, n=3 ($p<0.0147$).

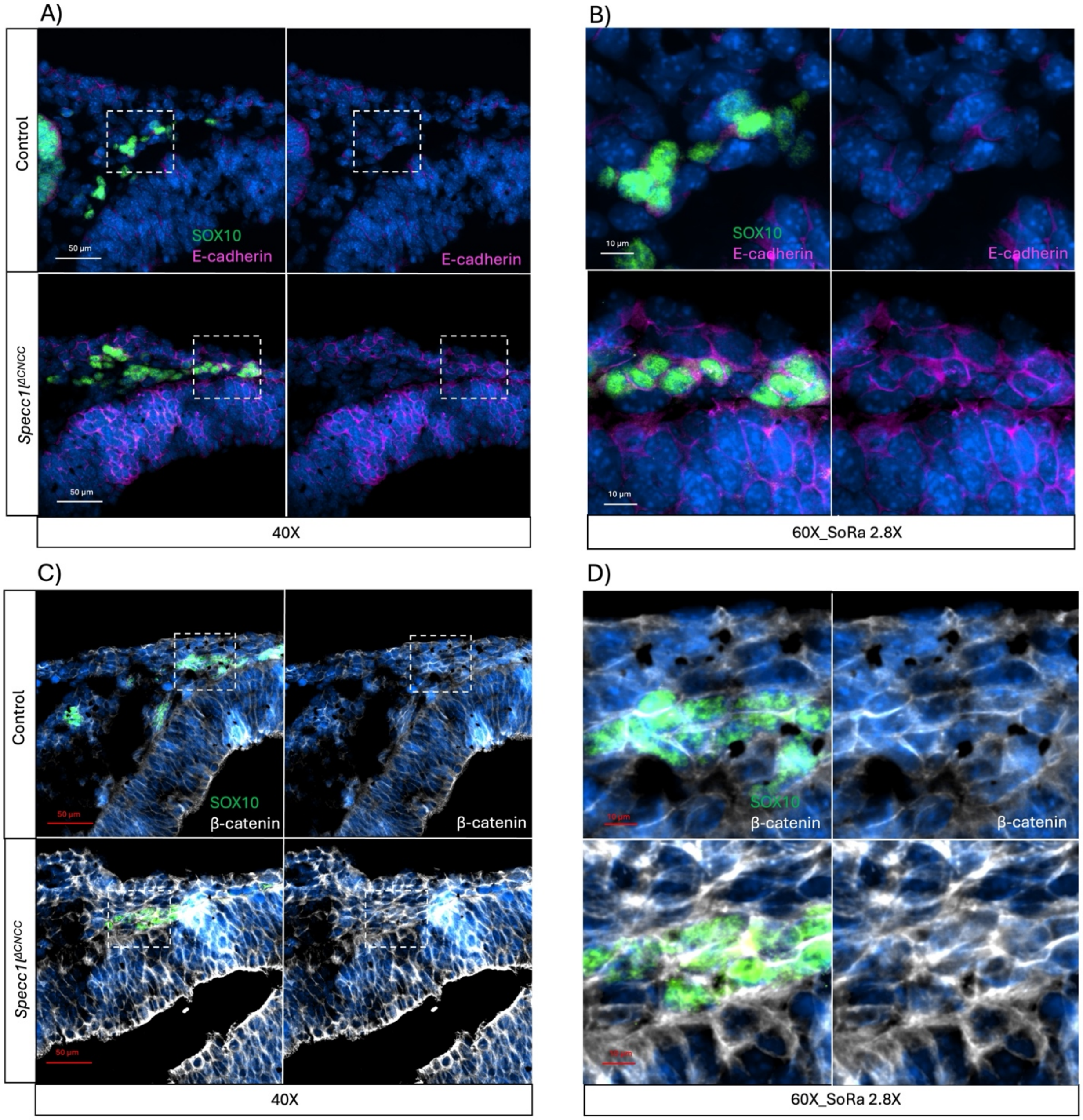


**Supplementary Figure 5:** Co-immunostaining of migratory neural crest marker SOX10 with adherens junction markers E-cadherin **(A, B)** or β-catenin **(C, D)** in control and mutant E9.5 embryos at 40x **(A, C)**. Boxed regions in A, C were magnified to 60x with SoRa 2.8x **(B, D)**. Both junctional markers showed increased ectopic staining in SOX10-positive cells.

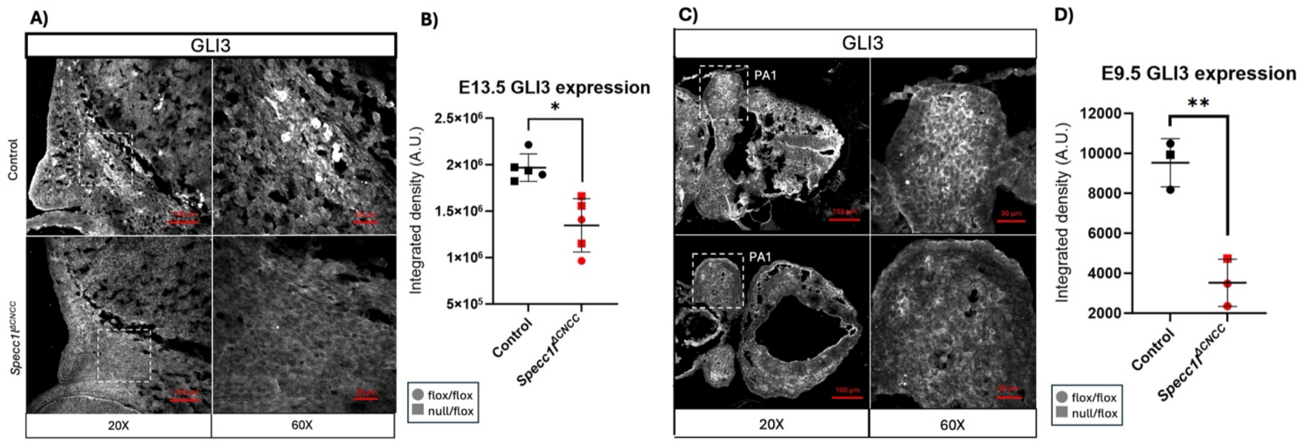


**Supplementary Figure 6:** GLI3 immunostaining at 20X and 60X magnification in E13.5 cranial mesenchyme **(A, B)** and in E9.5 first pharyngeal arch mesenchyme **(C, D).** GLI3 levels are decreased at both E13.5 **(B)** and at E9.5 **(D)**. Data represent mean ± SD. Statistical significance was assessed using an unpaired two-tailed *t*-test; n=5 for **B** and n= 3 for **D** (* P<0.05, ** P<0.01).
